# JASPER: a fast genome polishing tool that improves accuracy and creates population-specific reference genomes

**DOI:** 10.1101/2022.06.14.496115

**Authors:** Alina Guo, Steven L. Salzberg, Aleksey V. Zimin

## Abstract

Advances in long-read sequencing technologies have dramatically improved the contiguity and completeness of genome assemblies. Using the latest nanopore-based sequencers, we can generate enough data for the assembly of a human genome from a single flow cell. With the long-read data from these sequences, we can now routinely produce *de novo* genome assemblies in which half or more of a genome is contained in megabase-scale contigs. Assemblies produced from nanopore data alone, though, have relatively high error rates and can benefit from a process called polishing, in which more-accurate reads are used to correct errors in the consensus sequence. In this manuscript, we present a novel tool for genome polishing called JASPER (Jellyfish-based Assembly Sequence Polisher for Error Reduction). In contrast to other polishing methods, JASPER gains efficiency by avoiding the alignment of reads to the assembly. Instead, JASPER uses a database of k-mer counts that it creates from the reads to detect and correct errors in the consensus. In addition to its use for polishing genomes, JASPER can also create population-specific genomes using an existing reference genome along with sequencing reads from multiple individuals from the population of interest. In this mode, JASPER alters the reference genome so that it contains variants that are common in the target population. In our experiments, we show that after creating a Japanese-specific reference genome, we observed a 27% reduction in homozygous variant calls using whole-genome sequencing reads from an individual from Tokyo.

## Introduction

As a result of continual increases in sequencing efficiency and much longer read lengths, highly contiguous genome assemblies are now a staple of genomic research. In particular, instruments from Oxford Nanopore Technologies have enabled many scientists to sequence and assemble genomes from many species without the need for a significant investment in sequencing equipment. A single PromethION flowcell, for example, now yields enough data to produce a highly contiguous assembly of a human genome. However, because the per-base accuracy of ONT reads is still relatively low, creating an accurate final sequence (the “consensus”) remains a challenge. One way to improve the consensus quality is to “polish” the assembly using accurate reads from the same DNA source. These more-accurate reads can either be short (100-250bp) Illumina reads, or more expensive but longer (10Kb and above) PacBio HiFi reads. A variety of polishing tools have been developed and published for this purpose, including Pilon [1], Racon [2], POLCA [3], ntEdit [4], and NextPolish [5]. Almost all of these tools require aligning the polishing reads to the assembled contigs, which is computationally expensive. ntEdit does not require alignment, but instead uses Bloom filters to compute and store k-mer quality information about the reads used for polishing. In this manuscript, we introduce a novel alignment-free polishing tool, JASPER (Jellyfish-based Assembly Sequence Polisher for Error Reduction), which uses k-mer counts computed from the polishing reads to make corrections in assembled contigs. Our experiments show that JASPER is substantially faster than alignment-based polishing tools, with a relatively small cost in overall accuracy. We also show that JASPER is both faster and more accurate than ntEdit.

The computational efficiency of JASPER enables another possible application, which we illustrate in this study with an example. One can use JASPER to produce population-specific human reference genomes, using Illumina reads sequenced from multiple individuals from a population of interest. Aligning these Illumina reads to a genome could become quite expensive, especially if the number of individuals is large, while counting k-mers in the reads is much cheaper computationally. JASPER can use these k-mer counts to “correct” a human genome assembly so that it contains all homozygous variants that are common in the population from which the reads were drawn. When using this population-specific reference for comparisons of DNA from other individuals from the same population, shared variants will not be seen, thus making downstream analyses easier.

## Results

We first compare JASPER to several leading genome polishing tools, including POLCA [3], NextPolish [5], and ntEdit [4]. POLCA and NextPolish are among the best methods for polishing with Illumina reads [3], and ntEdit is the best of the alignment-free genome polishers. ntEdit is based on counting and classifying k-mers in a set of Illumina reads using Bloom filters, and then using the classification information to detect and fix errors in the consensus. In our first experiments, we utilized publicly available data for the model plant *Arabidopsis thaliana* (ecotype Col-0, see Data availability section). We began with a previously-published high-quality assembly of the genome [7], which we label Athal-Berlin, and introduced 39635 single base substitutions and 78928 single base insertions and deletions at random locations in the genome. We call the resulting assembly Athal-simerr. We then used wgsim (https://github.com/lh3/wgsim) to simulate 30x coverage by 2×150 paired-end Illumina reads with a 1% error rate. We call these reads the “simulated polishing reads.” We then used the simulated polishing reads to correct the Athal-simerr assembly with POLCA, NextPolish, ntEdit and JASPER. In the second experiment, we used the MaSuRCA assembler [8] (v4.0.6) to create another assembly of *A. thaliana* using the PacBio and Illumina reads from Berlin et al. [7]. We call this assembly Athal-MaSuRCA. We polished this assembly with POLCA, NextPolish, ntEdit, and JASPER, using a 60x coverage subset of the Illumina reads from Berlin et al. [7] with all methods. We used the default settings for polishing, with k-mer length = 25 for JASPER and ntEdit. For evaluation, we aligned the polished assemblies to the Athal-Berlin reference using MUMmer version 4 [9], and analyzed the alignments using the dnadiff tool from the MUMmer package. The dnadiff tool reports the number of insertion/deletion and substitution differences in the 1-to-1 best alignment of the two genomes. The reason why we prefer alignment-based validation as opposed to a k-mer-based quality assessment tool (e.g., Merqury [10]) is that spurious “corrections” in repeat regions can incorrectly make a copy of repeat look like the copy of the same repeat from a different place in the genome, wiping out small differences between the repeat copies. Such spurious corrections can reduce the number of incorrect k-mers in the assembly, resulting in an erroneous but higher assembly quality score. Alignment-based validation uses sequence information over longer spans, and ensures that repeats are aligned properly.

Our evaluation of the polishing tools on the *A. thaliana* data is shown in Tables 1 and 2. Table 1 shows that JASPER overall corrects more errors than ntEdit, but does not correct as many as the alignment-based polishing tools, POLCA and NextPolish. On the real data ntEdit actually increases the number of substitution errors slightly, while all methods reduce the number of indels. The primary reason for the inferior performance of the k-mer-based methods is that most differences between Athal-MaSuRCA and Athal-Berlin are indels of several bases in a row. While alignment-based methods are able to correct such errors, k-mer-based methods fail when the indel size is close to the size of the k-mer (25bp in these experiments). Table 2 shows more in-depth analysis of the specific corrections made by different polishing methods on the simulated data. The alignment-based methods perform best in this analysis, where POLCA is more accurate and NextPolish more sensitive. Here we see that JASPER has lower sensitivity compared to the alignment-based methods, but has excellent precision, bested only by POLCA. JASPER is superior to ntEdit in both sensitivity and precision.

**Table 1.** Comparison of polishing tools on assemblies of *A. thaliana* Col-0. Numbers correspond to the number of bases in substitutions, insertions, or deletions detected by aligning the polished assemblies to the corresponding reference, Athal-Berlin. Columns labeled “Simulated” refer to experiments that used a reference into which we introduced random errors, and that used short reads simulated from the unaltered reference for polishing. Columns labeled “Real” refer to experiments that used real Illumina reads to polish a draft assembly of the data. Timing data used a 32-core Intel Xeon server with 32 threads. Execution time is averaged across all 4 experiments. “Not corrected” refers to the number of single-base indels or substitutions that were not corrected after applying each tool. The best value in each column is shown in bold.

|  | Simulated errors<br>#bases in<br>substitutions<br>not corrected | Simulated errors<br>#bases in indels<br>not corrected | Real errors<br>#bases in<br>substitutions<br>not corrected | Real errors<br>#bases in indels<br>not corrected | Execution time<br>(minutes) |
| --- | --- | --- | --- | --- | --- |
| No polishing | 39635 | 78928 | 18295 | 29252 | n/a |
| POLCA | **76** | **234** | **16375** | 18494 | 121 |
| NextPolish | 721 | 1077 | 18054 | **18227** | 85.0 |
| ntEdit | 3992 | 5875 | 18367 | 27259 | 24.8 |
| JASPER | 954 | 4079 | 17827 | 27073 | **14.5** |

**Table 2.**
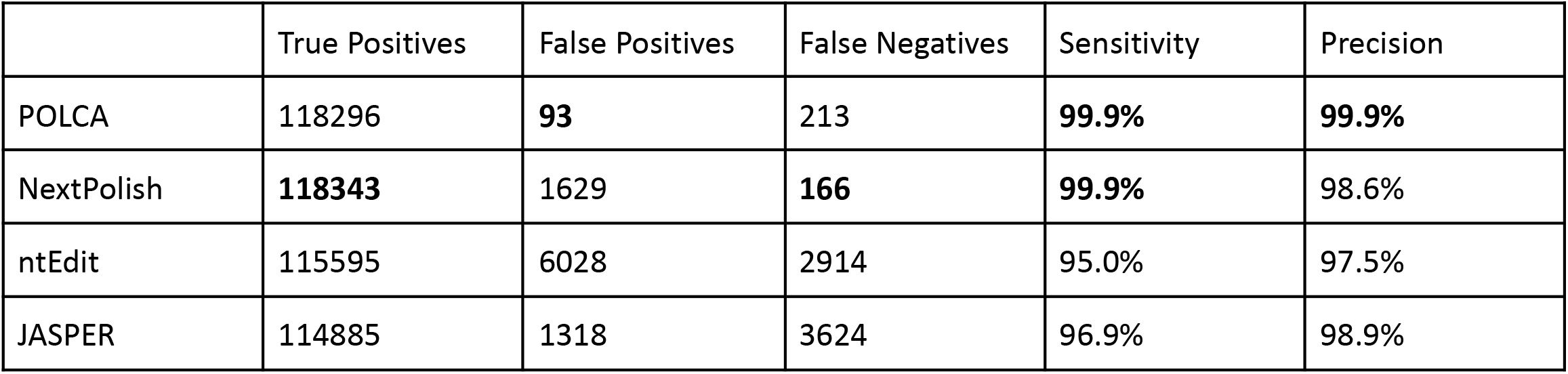
Statistical analysis of the corrections made by different polishing methods on an *A. thaliana* with simulated errors using simulated reads with no errors. True Positives are errors that we introduced and that were corrected; False Positives are spurious corrections that polishers made where there were no errors introduced; and False Negatives are errors in the genome that were not corrected.

|  | True Positives | False Positives | False Negatives | Sensitivity | Precision |
| --- | --- | --- | --- | --- | --- |
| POLCA | 118296 | **93** | 213 | **99.9%** | **99.9%** |
| NextPolish | **118343** | 1629 | **166** | **99.9%** | 98.6% |
| ntEdit | 115595 | 6028 | 2914 | 95.0% | 97.5% |
| JASPER | 114885 | 1318 | 3624 | 96.9% | 98.9% |

*A. thaliana* has a relatively small genome that lacks some of the complex repetitive structures that are present in mammalian genomes. To test our approach on a mammalian-sized genome, we used a public but unpublished assembly (GenBank accession GCA_001015355) of the human CHM13 cell line that was assembled with Celera Assembler v. 8.3rc2 from ∼70x coverage in PacBio P6C4 reads. Recently the T2T consortium published a very high quality assembly of the same CHM13 cell line, the first-ever truly complete human assembly [11]. This new CHM13 reference genome (v2.0) provides a straightforward way to compare polishing tools by using whole genome alignments to measure the number of substitution and indel differences between the GCA_001015355 assembly and CHM13 v2.0 before and after polishing.

We used nucmer [9] to align the original GCA_001015355 assembly and several polished versions of it to CHM13 v2.0, and then counted the number of differences between the assemblies. The un-polished GCA_001015355 assembly had 1,093,310 bases in indels and substitutions when compared to the CHM13 v2.0 reference. For polishing, we used 6,899,727 PacBio HiFi reads for CHM13 containing 75.6G bp of sequence (see Data Availability section). JASPER reduced the total number bases in consensus errors by 37,751, while ntEdit had a smaller reduction of 25,061. POLCA and NextPolish performed much better, reducing the number of errors by 135,810 and 138,275 respectively.

While alignment-based methods identify and correct somewhat more errors, JASPER was much faster, completing its polishing in ∼9 hours, followed by ntEdit which took ∼13 hours. In contrast, POLCA and NextPolish each took ∼97 hours, with the bulk of the time spent aligning the PacBio reads to the genome.

JASPER also reports an overall quality value (QV) for the assembly based on the fraction of erroneous k-mers it observes before and after correction. In this calculation, any k-mer in the assembly that does not appear in any read is considered an erroneous k-mer. We then compute the quality as QV = (−10)log_10_(E_b_), where E_b_ is the proportion of erroneous bases in the assembly. E_b_ is estimated from the observed number of erroneous k-mers as E_b_ = 1 - (1 - E_k_)^(1/K)^, where K is the k-mer length and E_k_ is the k-mer error rate. E_k_ is computed as the number of erroneous k-mers divided by the genome size; e.g., if we found 100 erroneous k-mers in a 1Mbp genome, then E_k_ would be 1/10000. This approximation assumes that the errors are distributed randomly in the genome, which is not entirely true; e.g., two or more errors might occur consecutively in homopolymers. We note that QV as computed here is an under-estimate of the true assembly quality, because it assumes that the reads contain all k-mers in the genome, which is generally not true. In real data sets, it is usually impossible to know how much of the assembled sequence is missing from the reads, especially if the genome was assembled with reads from one sequencing technology (e.g. Oxford nanopore) and polished with reads from another (e.g. Illumina), although that number is likely to be very small when coverage is deep. For example, even in our experiments with 30x coverage of *A. thaliana* using simulated 150bp reads, we found 1080 k-mers (out of about 120M) in the assembly that were not present in the simulated reads.

We can capitalize on the speed of JASPER to address another interesting problem: creating population-specific human reference genomes. When calling small variants (substitutions and indels) from whole-genome sequencing data of one individual against the reference genome, one typically gets several million variant calls. Some of these variant calls, especially those where the individual is homozygous, will be uninformative because they are common variants in the sub-population to which the individual belongs. To reduce the number of uninformative homozygous variant calls, one can use polishing to “correct” the reference genome in places where it disagrees with a majority of individuals from a given sub-population. Doing so will essentially hide the variants that are common in the sub-population to which the individual belongs, thus making downstream analysis easier. JASPER can create population-specific reference genomes by using Illumina data from multiple individuals to “polish” a human genome assembly such as GRCh38 or CHM13. In this application, we are not really polishing, but rather altering the assembly to make it look much more like a member of the target population. To illustrate this capability, we performed two experiments.

In the first experiment, we used the recently published complete CHM13 human genome [11] as the basis for creating a population-specific reference. Because CHM13 is missing the Y chromosome, we added the Y chromosome from GRCh38 to the CHM13 assembly, creating a new assembly which we designate CHM13+Y. We then downloaded Illumina data from eight individuals (NA18939, NA18946, NA18976, NA18978, NA18979, NA18981, NA18991, NA19011), all from Tokyo, Japan, from the 1000 Genomes Project website (http://www.internationalgenome.org). This yielded a total of ∼260 Gbp in Illumina reads. We then set aside one sample, NA18939, and used the remaining sets of reads to polish CHM13+Y using JASPER, resulting in an assembly we call Tokyo-polish.

We then aligned the reads from NA18939 to both the CHM13+Y and Tokyo-polish genomes, and called variants with POLCA [3]. We then counted the number of homozygous variant calls where the reference allele was supported by 0 or 1 reads (to allow for errors) and the alternate allele was supported by at least 3 reads. We found 1,590,668 such variants when comparing the reads to the CHM13+Y genome, and 1,161,050 variants when comparing to the Tokyo-polish reference. Thus, we observed a 27% reduction in the number of homozygous variant calls against the polished reference.

A related question is whether we would get a similar result if we called variants for an individual who was unrelated to the population of interest. To answer this question, we used Illumina reads for individual PGP17 from the Personal human Genome Project (PGP), who is estimated to be about 60% Ashkenazi with no Japanese ancestry. We observed 1,384,305 homozygous variant calls against the Tokyo-polish reference vs. 1,376,917 calls vs CHM13+Y. Thus (as expected) PGP17 was no closer to our Tokyo-polish assembly than to the CHM13+Y reference.

We then ran an experiment to compare the use of JASPER to create a population-specific reference versus using the recently-published Ashkenazi Jewish reference genome, Ash1 [14]. Ash1 was assembled from HG002, an Ashkenazi Jewish individual who is part of the 1000 Genomes Project. We also downloaded 30X coverage in Illumina data from his parents, HG003 and HG004. We used the Illumina reads from both parents to “polish” GRCh38.p13, creating a new Ashkenazi-specific reference, which we designated as GRCh38-Ash.

The publication describing Ash1 [14] called variants in the PGP17 genome by aligning the Illumina reads from PGP17 to Ash1, and found 1,333,345 homozygous variants, versus 1,700,364 homozygous variants when found when aligning PGP17 reads to GRCh38. We used the same methods and thresholds to call variants in PGP17 versus our GRCh38-Ash reference genome, and found 1,234,668 homozygous variants. Thus we found fewer homozygous variants, suggesting that our polished genome is an even closer match to PGP17 than the Ash1 genome. Note that this is not unexpected, because the reads used for polishing were from the two closest relatives of HG002.

In summary, JASPER is a very fast tool for polishing genomes, approximately 6 times faster than the fastest alignment-based polisher. In addition to its usefulness as a polishing tool, JASPER can also be used to efficiently create new personalized reference genomes.

## Methods

JASPER attempts to locate and correct errors in a genome sequence using only k-mer count data (k=25 by default) in a set of reads sequenced from the same individual. Given a set of reads, JASPER first calls Jellyfish to count k-mers in the reads. It then uses the “jellyfish histo” command to create a histogram of k-mer counts N(C), where N is the number of distinct k-mers with count C in the reads. The correction algorithm is implemented in Python, and it calls Jellyfish from within Python to load and query the database of k-mer counts. JASPER runs multiple instances of the correction algorithm in parallel, using shared memory to load the database.

Figure 1 shows a typical k-mer count histogram, with the k-mer count on the horizontal axis and the number of k-mers with a particular count on the vertical axis. The histogram is actually a combination of two different functions E(C) and R(C), where E(C) is the distribution of erroneous k-mers as a function of their count (shown in red in Figure 1) and R(C) is the number of true k-mers (from the genome) that occur in the sequence data (blue area in Figure 1). The local minimum in the histogram occurs at the point C_eq_ where E(C_eq_)∼=R(C_eq_), where any k-mer with a count of C_eq_ has a 50% chance of containing a sequencing error. K-mers with counts below C_eq_ are more likely to contain sequencing errors than to belong to the genome.

**Figure 1:**
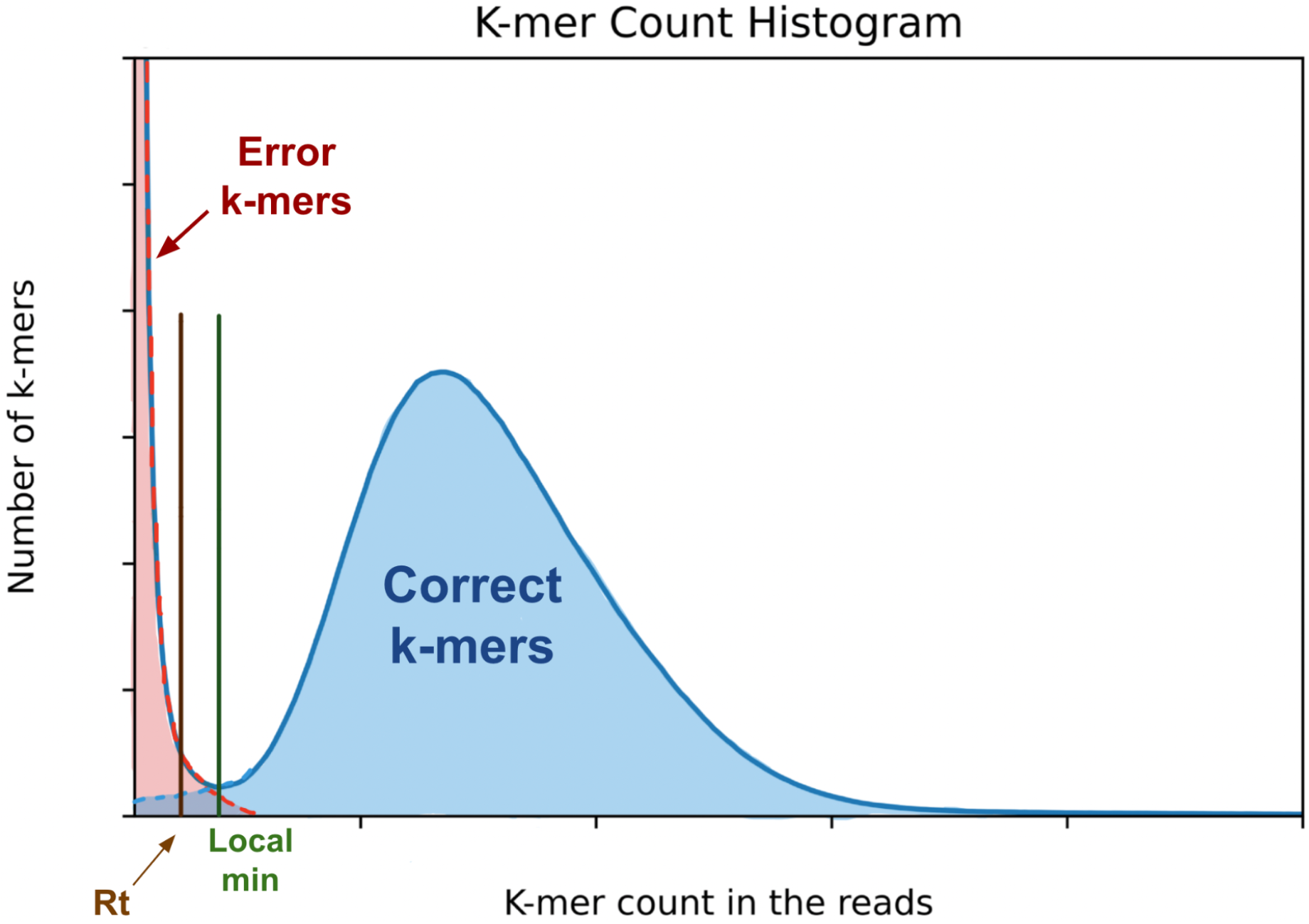
A typical k-mer count histogram for low-error-rate sequencing data (Illumina or PacBio HiFi). The red region contains error k-mers that are due to sequencing errors in the reads. The blue region represents the distribution of the counts of correct k-mers in the reads. The x-position labeled R_t_ is defined as half of the x coordinate of the local minimum of the distribution.

The general workflow of JASPER is as follows. For each contig sequence in an assembly, we look up the count for each k-mer in the contig in the database of k-mer counts (from the reads). We skip any k-mers containing a non-ACGT character. For each k-mer we determine whether it contains an error by using a heuristic approach with two thresholds: an absolute threshold, A_t_, and a relative threshold, R_t_, where A_t_ < R_t_ < C_eq_. Any k-mer with a count below A_t_ is considered to be erroneous. By default, JASPER sets R_t_=floor(0.5*C_eq_) and A_t_=floor(0.5*R_t_), where C_eq_ is the local minimum of the histogram curve as shown in Figure 1, and floor(x) is the closest integer to x that is smaller than x. We note that this computation works for C_eq_ >= 4. If C_eq_ < 4, then JASPER considers that the input read data is not suitable for polishing and returns an error. If a k-mer has a count less than R_t_ but greater than A_t_, then we label it as erroneous if its count is also less than half of the count of the previous k-mer. This heuristic is intended to account for natural variations of coverage in the sequencing data. As we examine k-mer counts along the genome sequence, any drop in k-mer counts that is due to natural variation in coverage is likely to be smooth, whereas any drop that is due to an error in the sequence is likely to be sharp. When we find the first erroneous k-mer K_e_ after a run of correct k-mers, all subsequent consecutive k-mers whose counts are below R_t_ are considered to be erroneous as well. JASPER continues to mark k-mers as erroneous until it sees a k-mer with count above R_t_. This generates a “run” of erroneous k-mers of length L. We use the length of the run L to determine the type of error in the contig sequence and the appropriate correction strategy, as follows:

1. L = K (25 by default): the putative error is a single-base substitution or insertion located at the last base of the sequence of errors, S_L_. JASPER attempts to fix this error by changing the last base of S_L_ or deleting it. We accept a fix if counts of all k-mers in the run are above R_t after the modification.
2. L > K: we assume that this is caused by two errors that are less than K apart. We assume that one or both errors are substitutions, and we attempt to fix them by modifying the last base of the first k-mer in S_L_, and the first base of the last k-mer in S_L_. We accept the full or partial fix if the counts of the first two k-mers and/or the last two k-mers in S_L_ are above R_t_ after the modification. This fix will fail if both errors are insertions or deletions, which is a limitation of our method. It will fully succeed if both errors are substitutions, and it will succeed partially if one of the errors is a substitution and the remaining indel may be fixed in a subsequent iteration of polishing.
3. L=K-1: this indicates that S_L_ contains a deletion immediately before the last base of the first bad k-mer. JASPER tries to fix it by inserting a base at that site. We accept the fix if all k-mers spanning the putative deletion site have counts at least R_t_ after the modification.
4. L<K-1, this could be caused by a substitution at the last base of the first k-mer in S_L_ or a substitution at the first base of the last k-mer in S_L_ (not both). This situation may occur because of a haplotype phasing error, where the assembly algorithm picked a base from one haplotype and a base from the other haplotype separated by K or fewer positions. The result is that all k-mers spanning the two erroneously phased bases may be invalid. Another scenario for this case to occur is an error in a repeat region, where the two bases that are closer than K apart come from two distinct copies of the repeat. This scenario is similar to haplotype phasing error. We try to fix this type of error by modifying one of the putative erroneous bases. We accept the fix if counts of all k-mers in S_L_ have counts greater or equal to R_t_. If L<K-1 and the previous fixes did not work, then the cause might be a homopolymer insertion or deletion.
5. An insertion error is only possible when the last base of the last good k-mer before S_L_ is the same as the last base of the first bad k-mer (in S_L_). JASPER tries to fix it by deleting the last base of the first bad k-mer. Repeat the deletion up to L times or until the last base of the last good k-mer before S_L_ is not equal to the last base of the first bad k-mer, or the number of error k-mers spanning the deleted base becomes zero. Reject the fix completely if at any iteration the number of error k-mers does not decrease. Accept the fix if the number of error k-mers spanning the deleted base becomes zero, that is all k-mers spanning the site of modification have counts above R_t_.
6. Otherwise, the putative error is likely to be a deletion in a homopolymer. JASPER will try to fix it by inserting the same base as the last base of the last good k-mer after the last base of the last good k-mer before S_L_. Repeat the insertion up to L times or until the number of error k-mers spanning the inserted bases becomes zero. Reject the fix completely if at any iteration the number of error k-mers does not decrease. Accept the fix if the number of error k-mers spanning the original deletion site becomes zero, that is k-mers spanning the modification site have counts above R_t_.

The major types of errors described above are illustrated on Figure 2. We use K=5 for clarity and space considerations. Note that for haplotype phasing error only one of the bases (last base of the first k-mer in S_L_, or first base of the last k-mer in S_L_) is wrong.

**Figure 2.**
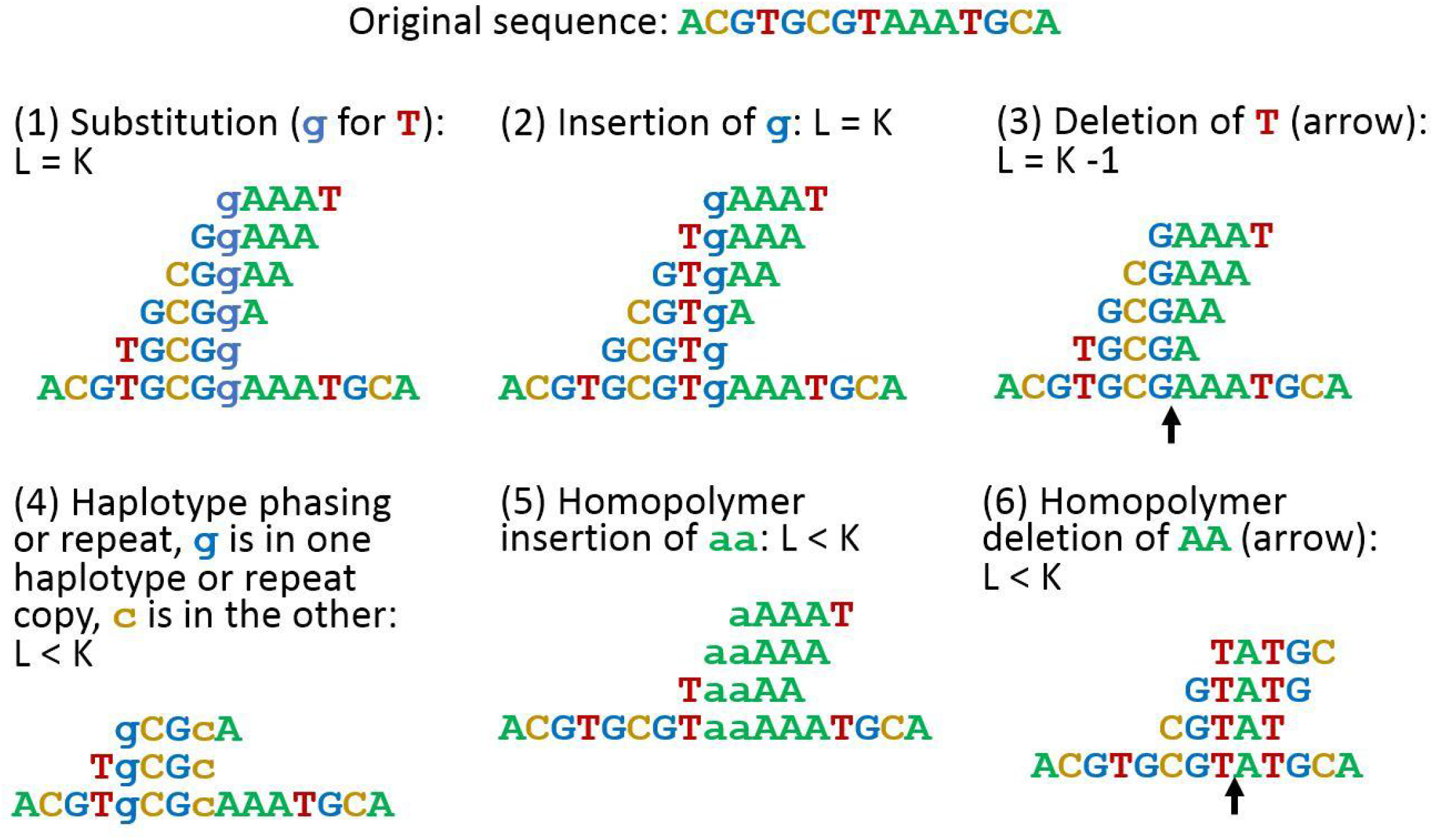
Illustration of the six error types that JASPER can detect and fix. Error bases are in lowercase. The error k-mers (K=5) are shown above the sequence. Black arrows indicate locations of deletions.

After we make a complete pass of corrections over the entire set of contig sequences, by default we run at least one more pass. Errors fixed on the first pass may allow JASPER to fix another set of errors on the second pass. For example, suppose the assembly contains two errors: a substitution and a deletion that are less than K bases apart (case 2, L>K). JASPER will only fix the substitution on the first pass. The deletion will become a standalone error, and JASPER will fix it on the second pass. Additional passes (which are optional) might correct more errors, although with rapidly diminishing benefits. Figure 3 shows the assembly QV value computed by JASPER and the number of corrections made in the *A. thaliana* simulated data experiment for 12 passes of polishing. The QV reaches its maximum after five passes, and the number of corrections made drops exponentially for the first five passes, and then flattens out completely at five corrections after the eighth pass.

**Figure 3.**
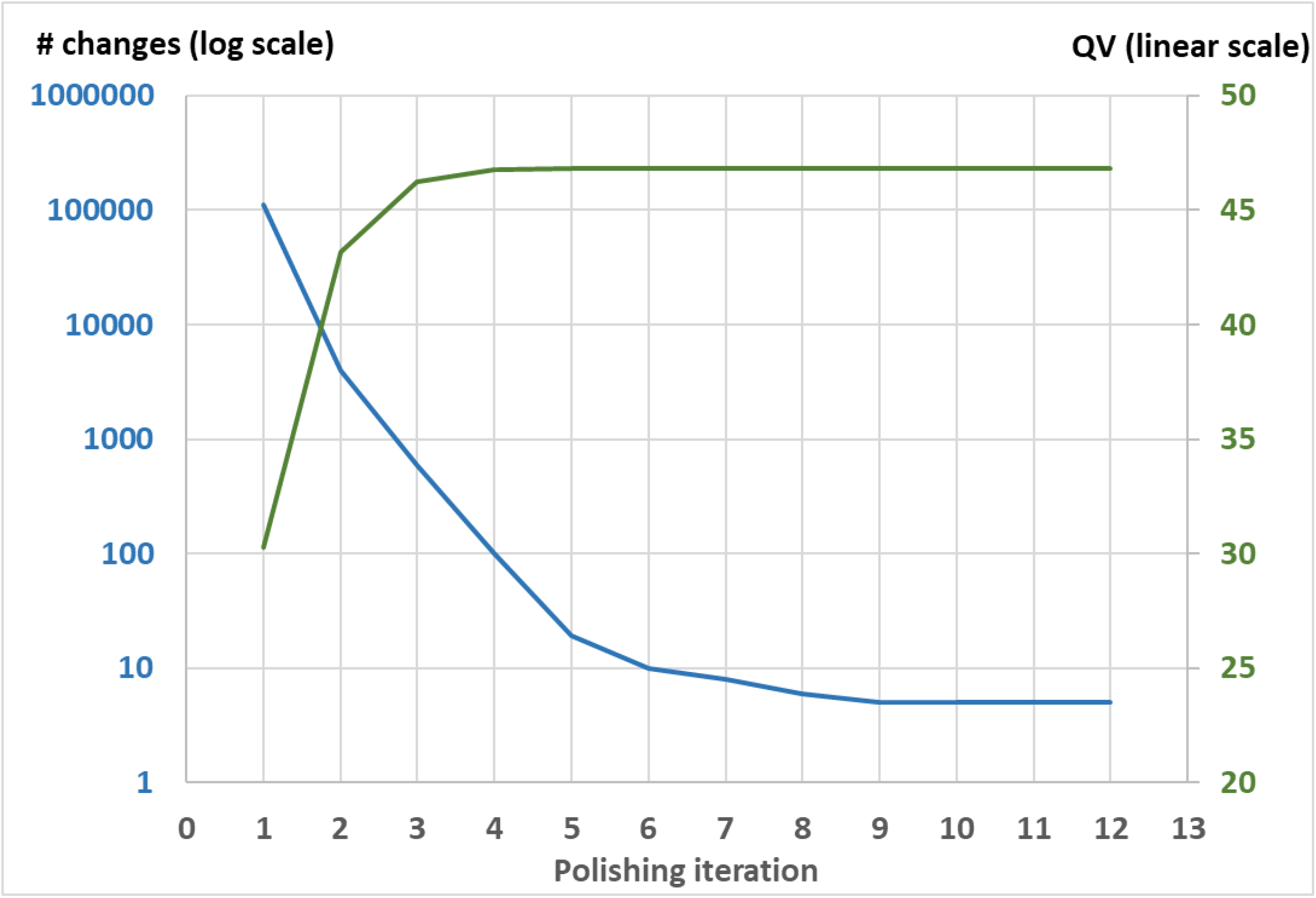
Assembly quality value (QV, in green, right axis, linear scale) and the number of bases changed by JASPER (in blue, left axis, log scale) in twelve successive iterations of polishing, on *A. thaliana* data with simulated errors. The QV depicted is measured before the polishing iteration; i.e. the QV at iteration 1 is the QV before any corrections were made.

## Summary

In this manuscript we introduced an efficient novel polishing tool called JASPER that is substantially faster than polishing methods based on sequence alignment, and more accurate than currently available k-mer based methods. The efficiency and scalability of JASPER allows one to use it to create personalized reference genomes for specific populations very efficiently, even for large sequenced populations. JASPER is open source software available on github at https://github.com/alguoo314/jasper.

## Data Availability

The Illumina and PacBio data for *A. thaliana* are available from NCBI SRA under accessions SRX533607, SRX533608 and from http://schatzlab.cshl.edu/data/ectools/. The human CHM13 data is available from https://github.com/marbl/CHM13 and from NCBI as Genbank accession GCA_009914755.4. The CHM13 assembly was polished with PacBio hifi reads from NCBI SRA accessions SRR9087597, SRR9087598,SRR9087599, and SRR9087600.

## References

1. Walker BJ, Abeel T, Shea T, Priest M, Abouelliel A, Sakthikumar S, Cuomo CA, Zeng Q, Wortman J, Young SK, Earl AM. pilon: an integrated tool for comprehensive microbial variant detection and genome assembly improvement. PloS ONE. 2014 Nov 19;9(11):e112963.

2. Vaser R, Sović I, Nagarajan N, Šikić M. Fast and accurate de novo genome assembly from long uncorrected reads. Genome Research. 2017 May 1;27(5):737–46.

3. Zimin AV, Salzberg SL. The genome polishing tool POLCA makes fast and accurate corrections in genome assemblies. PLoS computational biology. 2020 Jun 26;16(6):e1007981.

4. Warren RL, Coombe L, Mohamadi H, Zhang J, Jaquish B, Isabel N, Jones SJ, Bousquet J, Bohlmann J, Birol I. ntEdit: scalable genome sequence polishing. Bioinformatics. 2019 Nov 1;35(21):4430–2.

5. Hu J, Fan J, Sun Z, Liu S. NextPolish: a fast and efficient genome polishing tool for long read assembly. Bioinformatics (Oxford, England). 2019 Nov.Koren S, Walenz BP, Berlin K, Miller JR, Bergman NH, Phillippy AM. Canu: scalable and accurate long-read assembly via adaptive k-mer weighting and repeat separation. Genome Research. 2017 May 1;27(5):722–36.

6. Marçais G, Kingsford C. A fast, lock-free approach for efficient parallel counting of occurrences of k-mers. Bioinformatics. 2011 Mar 15;27(6):764–70.

7. Berlin K, Koren S, Chin CS, Drake JP, Landolin JM, Phillippy AM. Assembling large genomes with single-molecule sequencing and locality-sensitive hashing. Nature Biotechnology. 2015 Jun;33(6):623.

8. Zimin AV, Puiu D, Luo MC, Zhu T, Koren S, Marçais G, Yorke JA, Dvořák J, Salzberg SL. Hybrid assembly of the large and highly repetitive genome of Aegilops tauschii, a progenitor of bread wheat, with the MaSuRCA mega-reads algorithm. Genome Research. 2017 May 1;27(5):787–92.

9. Marçais G, Delcher AL, Phillippy AM, Coston R, Salzberg SL, Zimin A. MUMmer4: a fast and versatile genome alignment system. PLoS Computational Biology. 2018 Jan 26;14(1):e1005944.

10. Rhie A, Walenz BP, Koren S, Phillippy AM. Merqury: reference-free quality, completeness, and phasing assessment for genome assemblies. Genome biology. 2020 Dec;21(1):1–27.

11. Nurk S, Koren S, Rhie A, Rautiainen M, Bzikadze AV, Mikheenko A, Vollger MR, Altemose N, Uralsky L, Gershman A, Aganezov S. The complete sequence of a human genome. Science. 2022 Apr 1;376(6588):44–53.

12. Li H, Durbin R. Fast and accurate short read alignment with Burrows–Wheeler transform. Bioinformatics. 2009 Jul 15;25(14):1754–60.

13. Garrison E, Marth G. Haplotype-based variant detection from short-read sequencing. arXiv preprint arXiv:1207.3907. 2012 Jul 17.

14. Shumate A, Zimin AV, Sherman RM, Puiu D, Wagner JM, Olson ND, Pertea M, Salit ML, Zook JM, Salzberg SL. Assembly and annotation of an Ashkenazi human reference genome. Genome biology. 2020 Dec;21(1):1–8.

